# Longitudinal study of functional brain networks for processing infant directed and adult directed speech during the first year

**DOI:** 10.1101/2023.01.25.525490

**Authors:** Gábor P. Háden, Brigitta Tóth, István Winkler

**Affiliations:** Institute of Cognitive Neuroscience and Psychology, Research Centre for Natural Sciences, H-1117 Budapest, Magyar tudósok körútja 2., Hungary; Department Of Telecommunications And Media Informatics, Faculty of Electrical Engineering and Informatics, Budapest University of Technology and Economics, 1117 Budapest, Magyar tudósok körútja 2., Hungary

**Author notes:** Corresponding author: Gábor P. Háden, Institute of Cognitive Neuroscience and Psychology, Research Centre for Natural Sciences, Magyar tudósok körútja 2., Budapest H-1117, Hungary. Gábor P. Háden and Brigitta Tóth should be considered joint first author. **Author details** Brigitta Tóth, István Winkler. **Data Statement** Preprocessed data, analysis scripts and results will be made publicly available at the OSF repository. (DOI pending) https://osf.io/kd2tp/?view_only=5988023d9377426382210675b7170ac9. **Ethics** The study was conducted in full accordance with the Declaration of Helsinki and all applicable national laws, and it was approved by the relevant ethics committees: Medical Research Council-Committee of Scientific and Research Ethics (ETT-TUKEB), Hungary and United Ethical Review Committee for Research in Psychology (EPKEB), Hungary.

**Keywords:** development, listening, longitudinal, brain network topology, brain network topography, EEG, speech perception

## Abstract

In most cultures infant directed speech (IDS) is used to communicate with young children. The main role IDS plays in parent-child interactions appears to change over time from conveying emotion to facilitating language acquisition. There is EEG evidence for the discrimination of IDS form adult directed speech (ADS) at birth, however, less is known about the development of brain networks responsible for differentially processing IDS and ADS. The current study compared topological characteristics of functional brain networks obtained from 49 healthy infants at the age of 0, 6, and 9 months listening to the same fairy tale presented by the same speaker in IDS and ADS speech. Brain connectivity was assessed by the phase lag synchronization index in 6 frequency bands (delta, theta, low alpha, high alpha, beta, gamma). The topology of the large scale network organization was quantified using minimum spanning tree graphs, separately for each band. The delta band cortical network’s organization was found to be significantly more hierarchical and had a more cost-efficient organization during listening to ID compared to listening to AD. This network organization changes with age as nodes over the frontal cortex become more central within the network. The general picture emerging from the results is that with development the speech processing network becomes more integrated and its focus is shifting towards the left hemisphere. Our results suggest that ID speech specific differences in network topology are related to changes in the role of IDS during the first year of life.

**Highlights:**

- Multiple stages of maturation are reflected by different EEG bands, occurring in parallel, but with different timing.
- Networks processing infant directed speech changes during the first year of life reflecting the change in the role infant directed speech plays in development.
- Speech processing networks are shifting towards the left hemisphere with age.
- Longitudinal study of speech perception using functional networks on a large sample

## 1. Introduction

The course of language acquisition considerably varies across infants, and it is affected by how caregivers speak with their offspring (Ellis and Thal, 2008; Ramirez et al., 2020). In response to communicative stimulation by adults, neural networks distributed temporally and anatomically in infants’ brains develop/adapt to comprehend language (Friederici, 2002; Hickok & Poeppel, 2007; Meyer 2018). Infant directed speech (IDS) is a culturally near universal feature of adult to infant communication from birth (Fernald et al., 1989). There is evidence showing a distinct preference for IDS over adult directed speech (ADS) (e.g., Pegg et al 1992; Cooper et al., 1997; Fernald, 1985; Cooper & Aslin, 1990), which also changes throughout early development, a result that appears to be stable between multiple cultures and laboratories (ManyBabies Consortium, 2020). However, little is known about how and when this preference is represented in infants’ cortical networks. The aim of the present study was to longitudinally assess the organization and development of large-scale functional brain networks underlying speech processing in infants and furthermore to explore the change in sensitivity to infant directed speech. To this end, EEG was recorded from the same individuals listening to the same fairytale in both infant directed and adult directed speech registers at birth, at four months and at nine months of age.

### 1.1 The role of infant directed speech in development

Caregivers modify certain linguistic and prosodic aspects of their speech when they address infants (Fernald, 1985; Cooper & Aslin, 1990). The main linguistic features of IDS are fewer words per utterance, more repetitions, careful articulation, and decreased complexity (Fernald, et al., 1989; Cooper & Aslin, 1990). The prosodic IDS characteristics include exaggerated intonation, higher overall pitch, larger pitch changes, slower speech rate, longer pauses, and heightened emphatic stress (Cristia, 2013). Our previous study has shown that acoustic features of IDS are picked up by the brain already at birth (Háden, et al. 2020). Further, IDS helps infants to track speech in the cortical party situation (Kalashnikova et al., 2018).

There are at least three distinct but interrelated roles that IDS can play in development. These roles are not mutually exclusive and are hypothesized to become the focus of development at different ages. The first role, also the earliest in the course of development, is conveying emotional states (Trainor, Austin & Desjardins, 2000; Saint-Georges et al., 2013). Results showing larger frontal activations for IDS compared to ADS (Saito et al., 2007) in newborns and young (4-13 month old) children (Naoi et al., 2012) are compatible with the idea of emotional control. The second role of IDS is the facilitation of language learning, by facilitating speech segmentation, discrimination of phonemes (Thiessen et al., 2005). There is neural evidence for the IDS/ADS discrimination on such this level: infant directed vowels elicit larger responses over the right hemisphere (Zhang et al., 2011). Furthermore, discernible cues towards the grammatical structure of a language are also present in IDS (Fisher and Tokura, 1996; Soderstrom, 2007), although it is not clear whether and when these are utilized by the brain. Finally the third role that IDS may play in development is to direct attention towards the infant as the addressee of communication (Werker & McLeod, 1989; Senju & Csibra, 2008; Golnikoff et al., 2015). Specifically, the “ostensive signal hypothesis” points out that the intonation pattern of IDS serves as an ostensive signal, thus stimulating joint attention and helping affect sharing (Csibra, & Gergely, 2009; Csibra, 2010). Ostensive communication also appears as activity over central electrodes in 5 month olds (Parise and Csibra, 2013).

There is evidence suggesting that IDS has beneficial effects on language acquisition (Saint-Georges et al., 2013). Due to its effect on reading memory and mathematical skills, early language development is a predictor of the child’s future performance in educational systems. Therefore, it is important how caregivers speak with their offspring matters (Kalashnikova & Burnham, 2018). It is known that conversation-like interactions (such as the peekaboo game) have an effect on the baby’s brain activity and likely their brain development (Wass et al., 2020). Higher engagement in responsive parental communication during the first year of the infant’s life leads to better language skills of the infant (Shruti, Mastergeorge & Olswang, 2018). Conversely, one of the main indicators of the failure of response to the child’s needs is the “shallow” IDS pattern of depressed mothers (Herrera, Reissland and Shepherd, 2004).

### 1.2 The function of brain oscillations in speech processing

The development of speech comprehension and vocabulary learning require the skill of segmenting the continuous speech into meaningful units (Friederici 2002). At the age of seven month the infants are already capable of segmenting at least a few words reliably (Ma et al., 2011). Spoken language comprehension relies on temporally and anatomically distributed neural processes that extract the relevant phonological, grammatical, and lexical information (Friederici, 2002; Hickok & Poeppel, 2007; Luo & Poeppel, 2007; Meyer 2018). Neural oscillations are suggested to play a crucial role during speech parsing by defining the temporal boundaries between linguistic items within the continuous acoustic signal [such as the phonemic (∼25 ms or ∼40 Hz), syllabic (∼200 ms or ∼5 Hz), or phrase/sentence/context level (> 1000 ms or < 1 Hz; Giraud & Poeppel, 2012)]. Indeed, neural oscillations entrain (synchronize) to the syllabic rate of speech in a sustained manner, and neural entrainment to ongoing speech is dependent on the rate of preceding speech. Importantly, brain-to-speech synchrony at specific rates directly affects how words are understood (Dikker et al., 2019). The neural processes generating low-frequency delta and theta activity in auditory cortex were found to be synchronized with the speech envelope, suggesting that these oscillations parse the sound input at these timescales (Ghitza, 2011; Doelling et al 2014; Rimmele et al., 2015; Keitel et al., 2017; Luo & Poeppel, 2007).

During speech perception widely specialized distributed cortical networks are dynamically engaged in parallel for accumulating information over multiple timescales (Hill & Miller 2009; Salmi et al., 2009; Saur et al., 2008, 2010; Scott et al. 2009, 2013; Watkins et al., 2007). Long-range oscillatory synchronization (also termed functional connectivity – FC) is a potential candidate for assessing the operation of such neural mechanisms, as it measures coupling between cortical regions as a function of time (for reviews, see Buzsáki & Draguhn, 2004; Sporns, 2011). For example, Obleser and colleagues (2007) found that the success of comprehension of spectrally degraded speech is correlated with FC strength between cortical areas outside auditory cortex (angular gyrus, medial and lateral prefrontal cortices, posterior cingulate gyrus), assumed to be involved in higher-order processes of speech comprehension.

### 1.3 Brain network analysis and brain network development during childhood

The spontaneous dynamic interaction between distributed networks of functionally specialized brain areas changes across the early lifespan (from birth till four months) with important consequences for cognitive development through the neuronal mechanisms of learning and plasticity (Sporns, 2011). The organization of these functionally connected brain areas can be examined by brain networks represented as graphs, an abstract mathematical description of the elements of the network and their interactions (for reviews, see Bassett & Sporns 2017; Stam, 2014; Varela et al., 2001). In healthy adults EEG/MEG functional networks exhibit 1) an economic trade-off between high level clustering and short path (“small-world” organization, which provides optimal ratio of direct communication between closely spaced and separated brain areas; Stam & van Straaten, 2012), 2) hierarchical modular topology (De Haan et al., 2012; Van den Heuvel & Pol, 2010), and 3) include densely connected “hub” regions, which can serve as coordination centers (Achard et al., 2006).

This efficacy of communication between brain regions during rest is assumed to be crucial for healthy cognitive function (Bassett & Bullmore 2009; Van den Heuvel et al., 2009), and it is relevant for the development of cortical circuits as demonstrated by the involvement of neural synchrony in synaptic plasticity and changes in the synchronization of neural oscillations during development (Uhlhaas et al., 2009, 2010). Generally functional connections develop throughout the human lifespan (Fransson et al., 2007; Uhlhaas et al., 2009, 2010), and the development of synchronized oscillations between frontal and parietal regions and between the hemispheres is crucial for cerebral maturation (Grieve et al., 2008, Homae et al., 2010).

Only a few studies assessed the early development of brain networks measured with EEG in healthy newborns (Omidvarnia et al., 2014; Tokariev et al., 2015; Toth et al., 2017) showed network reorganization resulting from maturation or change in the external stimulation after birth. For example, brain maturation processes may play an important role in the relationship between infantile joint attention and language development (Mundy et al 2003). Subnetworks further develop later during infancy and they are gradually differentiated during childhood, probably linked with the development of cognitive abilities (Ferreira & Busatto, 2013; Van den Heuvel & Pol, 2010). Abnormal topology of the FC map has been suggested to underlie a wide range of neurocognitive brain disorders, including autism spectrum disorders (Uhlhaas & Singer, 2007). Thus slower maturation of FC during early postnatal period may predict subsequent delay in language acquisition.

### 1.4 Study aims

The present longitudinal study followed infants from birth to 9 months recording the electroencephalogram (EEG) while the infants were presented with ID and AD speech. We assumed that important aspects of the developmental trajectory of speech processing during the first year of life can be extracted from the EEG brain networks: 1) the development of speech processing brain networks at different timescales, and 2) changes in the sensitivity to IDS relative to ADS.

## 2 Materials and methods

### 2.1 Participants

EEG was recorded from 49 healthy, full-term infants (21 male) three times at the ages of 1-4 days (0 months), 120-151 days (4 months), and 274-317 days (9 months) postpartum. All infants were firstborns, none of them twins. The mean gestation age was 39.95 weeks (SD=0.98), the birth weight 3411 g (SD=354.96). Data recorded from 11 infants were excluded based on the criterion of retaining 40.96 seconds of EEG signal after artifact rejection (at least 10 epochs of 4096 ms duration) in all experimental conditions at all ages. Thus, 38 infants (14 male) were included in the final sample. (See details in Supplement 1). Informed consent was obtained from one (mother) or both parents when the family was recruited for the longitudinal study. The study was conducted in full accordance with the Declaration of Helsinki and all applicable national laws, and it was approved by the relevant ethics committees: Medical Research Council-Committee of Scientific and Research Ethics (ETT-TUKEB), Hungary and United Ethical Review Committee for Research in Psychology (EPKEB), Hungary.

### 2.2 Stimuli and procedure

#### 2.2.1 Procedure at birth

Infants were presented with recordings of a fairy tale delivered in Hungarian in both IDS and ADS register by a native Hungarian-speaking mother, who at the time of recording was directing her words to her own 4.5 month old infant or to the experimenter, respectively. Recordings were carried out in an anechoic room using a single sound channel and were later converted to dual-mono.

The durations of the two recordings differed from each other (ADS=104 s; ISD=144 s). Recordings were presented binaurally using the E-Prime stimulus presentation software (Psychology Software Tools, Inc., Pittsburgh, PA, USA) with ER-1 headphones (Etymotic Research Inc., Elk Grove Village, IL, USA) connected via sound tubes to self-adhesive ear cups (Sanibel Supply, Middelfart, Denmark) placed over the infants’ ears. Stimuli were presented in two blocks. Within each block, the recording of each speech register was presented two times in a row, with an inter-stimulus interval of 10 s between recordings within the blocks (overall eight stimuli, 4 presentation of each recording, ca. 18 minutes duration). The order of the speech registers was counterbalanced between the blocks and randomized across infants.

The experiment took place at the Department of Obstetrics-Gynecology and Perinatal Intensive Care Unit, Military Hospital, Budapest, with only the infant and the experimenter being present in the room. This experiment was part of a session combining multiple experiments, which were always presented in the same order: 1) fairy tale in IDS and ADS (Block I. of the current study), 2) Hungarian words delivered in IDS and ADS (5 min., same speaker as in the current study; data published in Háden et al., 2020), 3) entrainment to speech-like artificial sounds (15 min.), 4) an oddball sequence with tones of two different durations (15 min.); 5) fairy tale in IDS and ADS (Block II. of the current study), and 6) resting state (no stimulation; 5 min). The session was part of a longitudinal study aimed to assess the role of IDS and ADS in speech acquisition. Data from the other experiments and the associations between them will be published separately after collecting the complete data set.

#### 2.2.2 Procedure at four and nine months of age

The stimuli were identical to the newborn age recordings, however the very limited time available in these age groups necessitated to only present one stimulus/register (ca. 4.5 minutes overall duration). Stimuli were presented in a sound attenuated room using Matlab (MathWorks Inc., Natick, MA, USA) and Psychtoolbox (Kleiner, Brainard and Pelli, 2007). The sound signal from the computer was amplified by a Yamaha A-S301 amplifier (Hamamatsu, Japan) and presented through a pair of speakers at a comfortable intensity (Boston Acoustics A25, Woburn, MA, USA). The speakers were positioned ca. 1.75 meter in front of the participant, 70 cm apart.

This experiment was part of a session combining multiple experiments. As it was always the first experiment, the rest of the experiments are not listed. The experiment took place at the sound-attenuated and Faraday-shielded infant EEG laboratory of the Institute of Cognitive Neuroscience, Research Centre for Natural Sciences. The infant sat comfortably in his/her mother’s lap while the experimenter employed toys to keep the infant facing towards the loudspeakers and his/her attention away from the electrode net. The mother was listening to music through closed can audiometric headphones to isolate her from the experimental stimulation. If the infant became fussy, the playback stopped and the experimenter attempted to pacify him/her. If pacifying was successful the playback restarted from the beginning, otherwise the experiment was discontinued.

### 2.3 EEG recording

#### 2.3.1 EEG recording at birth

EEG was recorded during quiet sleep with Ag/AgCl electrodes attached to the scalp at the Fp1, Fp2, Fz, F3, F4, F7, F8, T3, T4, Cz, C3, C4, Pz, P3, P4 locations (location naming is in accordance with the international 10-20 system). The reference electrode was placed on the tip of the nose and the ground electrode on the forehead. EEG was digitized with 24 bit resolution at a sampling rate of 1 kHz by a direct-coupled amplifier (V-Amp, Brain Products GmbH, Germany). The signals were on-line low-pass filtered at 110 Hz. The impedance values were monitored throughout the recording session and were kept under 10 kΩ where possible, but no recordings were rejected based on impedance measurements. Although high impedance can introduce additional noise into the recordings, it may not be systematic in the entire group and its effects are mitigated by filtering out the very low frequencies, which is more likely to be contaminated by drifts. Infants’ sleep state was determined based on behavioral criteria (Anders, Emde & Parmelee, 1971). Infants were predominantly asleep during the recordings, in quiet sleep 82% of the time, and in active sleep 8.2% of the time.

#### 2.3.2 EEG recording at four and nine months of age

EEG was recorded using a 60-channel HydroCel GSN net (64 channel v1.0 layout, channels 61-64 are connected to ground in the small pediatric caps used here) with an GES 400 DC amplifier passing the digitized signal to the NETSTATION v5.4.1.1 software (both Electrical Geodesics, Eugene, OR, USA). Signals were recorded online at a sampling rate of 1000 Hz with the Cz reference. Electrode impedance during recording was maintained below 50 kΩ. Electrodes corresponding to the Fp1, Fp2, Fz, F3, F4, F7, F8, T3, T4, Cz, C3, C4, Pz, P3, P4 locations were selected for further analysis in order to keep the data compatible with the recordings in newborns.

### 2.4 EEG data analysis

EEG was analyzed with the EEGLAB (Delorme and Makeig, 2004) open source Matlab (MathWorks Inc., Natick, MA, USA) toolbox. The signals were first bandpass filtered (using zero phase band-bass, Hamming windowed sinc FIR filter; filter order 33000) to the 0.5-45 Hz frequency range, then re-referenced to average of all electrodes. Continuous EEG data were segmented into 4096 ms long epochs with 50% overlap between successive epochs. This epoch length is sufficient to assess oscillatory activity in the delta-to-beta frequency bands since it covers at least two cycles of the highest frequency (Fraschini, et al. 2016; Hildebrand, et al. 2012; Omidvarnia et al. 2014). Segments contaminated with physiological (eye movements, muscle artefacts) or external (i.e. environmental noise) artefacts were rejected by visual inspection. A minimum of 10 epochs was considered sufficient (Hillebrand et al., 2012; Tewarie et al., 2015) for further functional connectivity analysis. The final dataset consisted of an average of 49.81 (SD = 17.13) epochs/participant. Epochs were then filtered (using zero phase band-bass, Hamming windowed FFT; filter order 1650) into five frequency bands (delta: (0.5)-4 Hz; theta: 4-8 Hz; alpha: 8-12 Hz; beta: 12-30 Hz; gamma: 30-(45) Hz;).

The strength of FC was calculated as pairwise phase synchronization strength between pairs of EEG electrodes, separately for the five frequency bands and for each epoch. The phase synchronization was measured by the PLI index (developed by Stam et al., 2007). PLI was calculated by using the BrainWave software version 0.9.151.5 (available at http://home.kpn.nl/stam7883/brainwave.html). PLI is an undirected measure of connectivity that calculates the consistency by which one signal is phase leading or lagging with respect to another signal.

### 2.5 Graph-theoretical measures

Network analysis was performed on the graph theoretical representation of the functional connectivity matrix using the so-called minimum spanning tree (MST) approach (Figure 1A; for reviews, see Boersma et al., 2013; Stam & van Straaten, 2012).The MST approach has been successfully employed for describing FC network properties (e.g. hierarchical structure, degree distribution etc.) of healthy newborn infants (Tóth et al., 2017). In the current study, the edge-weighted undirected graph consists of nodes that are the EEG electrodes (N=16) and edges (N=(16×16)/2-16) that are the connectivity values between each pair of nodes and the edge weights are the inverse PLI. MST is a subset of the edges (N=16) of a graph that connects all the nodes together without cycles (creating only one possible path between each pair of nodes) and with the minimum possible total edge weight (using the edges with the strongest FCs). MST networks were constructed by Kruskal’s algorithm (Kruskal, 1956). MST was constructed separately for each epoch and frequency bands.

**Figure 1.**
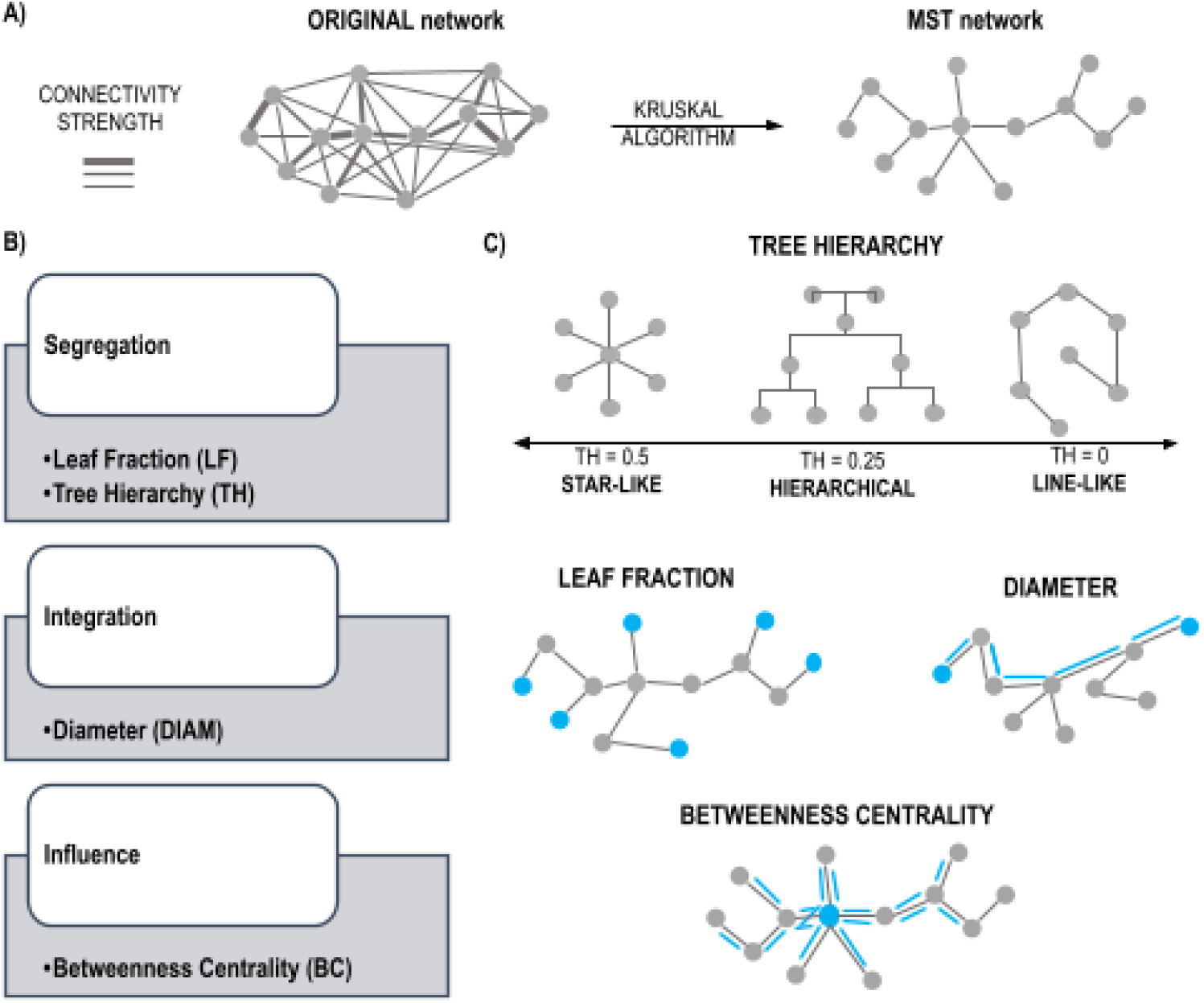
List of brain network measures used in the study. **A)** Schematic representation of network construction method. **B)** List of measures according to network property categories. **C)** Schematic illustration of MST measures.

Graph measures described by Stam and colleagues (2014; see also Tewarie et al., 2015) are illustrated on Figure 1B and C. MST network characteristics values were normalized by dividing them by the number of EEG channels used. The global MST network characteristics were averaged across epochs, separately for each infant and frequency band. The level of integration of the communication within the network was assessed by the metric called “Diameter” (DIAM). DIAM is the largest distance between any two nodes within the MST, where distance refers to the minimum number of edges required to proceed from one node to another (the shortest path). The degree of segregation within a network is measured by the metric of “Leaf Fraction” (LF) and “Tree Hierarchy” (TH). Leaf Fraction” (LF) is the number of nodes with only 1 connected edge divided by the total number of nodes in the MST. “Tree Hierarchy” (TH) assesses how hierarchical a given network is compared to the so-called ‘star-like network organization’ (for a mathematical description, see Boersma et al., 2013 and Tewarie et al., 2015). TH ranges from 0 (indicating a line-like topology) to 1; for the star-like topology, TH approaches 0.5. The level of influence or centrality of the nodes within the network elements was assessed by the metric “Betweenness Centrality” (BC). BC is a measure of the node’s ‘hubness’ within the network. It is defined as the normalized fraction of all shortest paths connecting two nodes that pass through the particular node (Newman & Girvan, 2010; Stam et al., 2014). BC is calculated separately for each node. BC may also be used to identify regions of the brain that serve as hubs by plotting the distribution of BC over the scalp (see, e.g., Tóth et al., 2017).

### 2.6 Statistical analysis

Statistical analysis was performed with the STATISTICA software package. Greenhouse-Geisser correction was employed to correct for sphericity violations (Greenhouse & Geisser, 1959) where appropriate and correction factor (ε) is reported where appropriate, as well as η_p_^2^ effect size value. Tukey correction was applied to the post-hoc pairwise comparisons. Both topological (LF, DIAM, and TH) and the topographical (BC) measures were used to characterize the development of processing ID and AD speech during the first year of life.

#### 2.6.1 Topology of the brain networks

For testing whether the brain networks affected by development (AGE, levels of 0 vs. 4 vs. 9 month) and by the SPEECH REGISTER (with level of ID vs. AD), repeated-measures ANOVAs were used to compare the characteristics of the infant FC networks, separately for each frequency band and network measure (LF, DIAM, and TH). Due to the potential confound caused by the different recording setup and sleep/awake state between newborns and the other two measurement times, we performed a secondary analysis testing network reorganization from four months to nine months of age by limiting the above described ANOVAs to 4 vs. 9 month of age.

#### 2.6.2 Topography of the brain network hubs

For testing whether the hub nodes are affected by development during the first year, repeated-measure ANOVAs were conducted for BC, separately for each frequency band by using AGE (0 vs. 4 vs. 9 month), SPEECH REGISTER (ID vs AD) and ELECTRODE (the 15 electrodes common to all three age groups). Similarly to the topological variables, the same analysis was repeated with only the 4-vs. 9-month old infants’ data included.

## 3 Results

Consistent results were only obtained for the delta, theta, and beta bands. These will be described in the main text. For all five frequency bands, all topological variables and the BC values for all electrodes can be downloaded from <anonymized osf.io link> (under Results).

### 3.1 Development of speech processing networks during the first year of life

Significant statistical results for the ANOVAs including all three groups are given in Table 1, while those comparing only the 4-month and 9-month old infants in Table 2.

**Table 1.** All three age groups: Summary of the significant effects involving AGE on the topological (upper table half) brain network characteristics (rows), separately for each frequency band with significant effects (columns). Significant post-hoc pair-wise test results, where applicable, are shown under the ANOVA in the following format: “Age in months” [0/4/9m] [</>] “Age in months” [0/4/9m] (p value). Post-hoc ANOVAs for the electrodes with significant age effects on “Betweennes Centrality” (BC) are shown in the lower half of the table.

| CHARACTERISTIC | EFFECT | DELTA BAND | THETA BAND | BETA BAND |
| --- | --- | --- | --- | --- |
| TH | AGE | F(2,74)=7.638, p=0.001<br>$\epsilon=0.997$, $\eta^2=0.171$;<br>0m < 9m (p=0.015), 0m < 4m (p<0.001) | | |
| DIAM | AGE | F(2,74)=16.190, p<0.001,<br>$\epsilon=0.951$, $\eta^2=0.304$;<br>0m > 9m (p<0.001), 0m > 4m (p<0.001) | F(2,74)=7.547, p=0.002,<br>$\epsilon=0.880$, $\eta^2=0.169$;<br>0m > 9m (p=0.003); 0m > 4m (p=0.004) | |
| LF | AGE | F(2,74)=16.574, p<0.001,<br>$\epsilon=0.990$, $\eta^2=0.309$;<br>0m < 9m (p<0.001), 0m < 4m (p<0.001) | F(2,74)=3.909, p=0.026,<br>$\epsilon=0.961$, $\eta^2=0.096$;<br>0m < 9m (p=0.029), 0m < 4m (p=0.085; tendency!) | |
| BC | AGE | F(2,74)=18.633, p<0.001,<br>$\epsilon=0.973$, $\eta^2=0.335$;<br>0m>4m (p<0.001), 0m>9m (p<0.001) | F(2,74)=8.044, p=0.001,<br>$\epsilon=0.922$, $\eta^2=0.179$;<br>0m>4m (p<0.004), 0m>9m (p<0.002) | F(2,74)=3.292, p=0.048,<br>$\epsilon=0.899$, $\eta^2=0.082$;<br>0m>4m (p<0.086; tendency!), 0m>9m (p<0.066; tendency!) |
| | AGE × ELECTRODE | | F(28,1036)=1.816,<br>p=0.036, $\epsilon=0.480$,<br>$\eta^2=0.047$ | F(28,1036)=2.199,<br>p=0.008, $\epsilon=0.472$,<br>$\eta^2=0.056$ |
| ELECTRODE-WISE POST HOC TESTS FOR THE ABOVE INTERACTION ON BC |  |  |  |  |
| ELECTRODE | INTERACTION |  |  |  |
| Fz | AGE | | F(2,74)=8.398, p=0.001,<br>$\epsilon=0.996$, $\eta^2=0.185$;<br>0m > 4m (p=0.008). 4m > 9m (p=0.001) | |
| C3 | | | F(2,74)=4.246, p=0.022,<br>$\epsilon=0.896$, $\eta^2=0.103$;<br>0m > 9m (p=0.021) | |
| Pz | | | F(2,74)=6.136, p=0.007,<br>$\epsilon=0.794$, $\eta^2=0.142$;<br>0m < 9m (p=0.003) | |
| P3 | | | F(2,74)=3.766, p=0.04,<br>$\epsilon=0.766$, $\eta^2=0.092$;<br>0m > 9m (p=0.022) | F(2,74)=4.504, p=0.022,<br>$\epsilon=0.793$, $\eta^2=0.109$;<br>0m > 4m (p=0.016) |
| Fp1 | | | | F(2,74)=3.663, p=0.031,<br>$\epsilon=0.990$, $\eta^2=0.09$;<br>0m < 9m (p=0.03) |
| Fp2 | | | | F(2,74)=6.238, p=0.004,<br>$\epsilon=0.963$, $\eta^2=0.144$;<br>0m > 9m (p=0.023), 4m > 9m (p=0.004) |

**Table 2.** Four and nine month olds only: Summary of the significant effects involving AGE on the topological (upper table half) brain network characteristics (rows), separately for each frequency band with significant effects (columns). Significant post-hoc pair-wise test results, where applicable, are shown under the ANOVA in the following format: “Age in months” [4/9m] [</>] “Age” (p value). Post-hoc ANOVAs for the electrodes with significant age effects on “Betweennes Centrality” (BC) are shown in the lower half of the table.

| CHARACTERISTIC | EFFECT | BETA BAND |
| --- | --- | --- |
| BC | AGE × ELECTRODE | F(14,518)=3.303 p=0.001 $\epsilon=0.550$<br>$\eta^2=0.082$ |
| ELECTRODE-WISE POST HOC TESTS FOR THE ABOVE INTERACTION ON BC |  |  |
| ELECTRODE | INTERACTION |  |
| Fp2 | AGE | F(1,37)=12.037, p=0.001, $\eta^2=0.245$;<br>4m > 9m (p=0.001) |
| F7 | | F(1,37)=4.325, p=0.045, $\eta^2=0.105$;<br>4m < 9m (p=0.045) |
| T3 | | F(1,37)=7.043, p=0.012, $\eta^2=0.160$;<br>4m < 9m (p=0.012) |
| Pz | | F(1,37)=4.430, p=0.042, $\eta p^2=0.107$;<br>4m < 9m (p=0.042) |

#### 3.1.1 Brain network topology

Characteristic differences were found between newborns and infants in the two later age groups (see Figure 2). The delta and theta band network structure became more hierarchical, star-like (increased Tree Hierarchy) with shorter paths to transfer information between nodes (shorter DIAM) and consisting of more peripheral nodes (higher LF) from 0 to 4 and 9 months of age.

**Figure 2.**
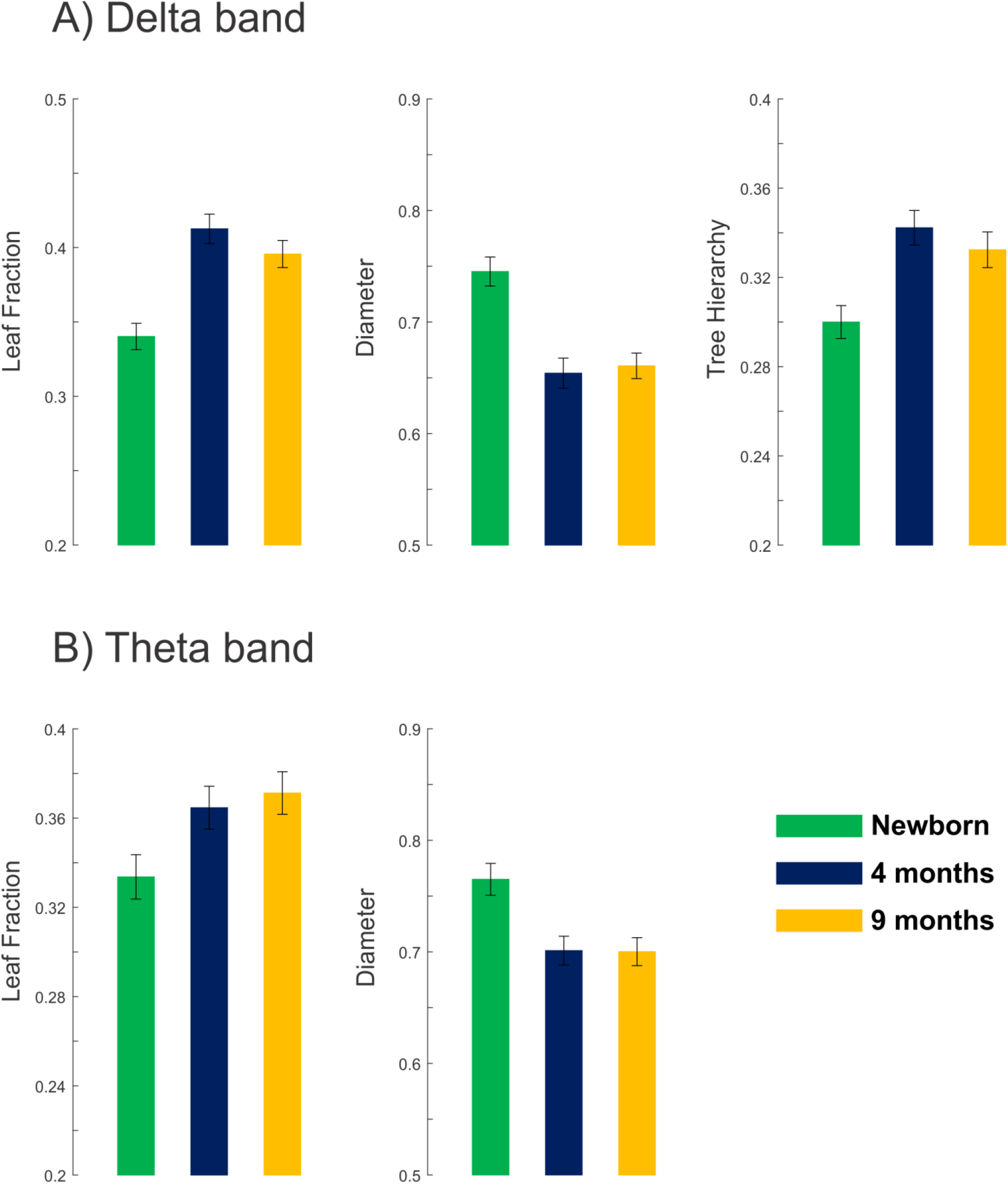
Development of topological characteristics of brain networks in the delta (panel A) and theta (panel B) bands. Error bars denote standard error of the mean (SEM).

When only considering the ages of 4 and 9 months, there were no significant effects of AGE on LF, TH and DIAM.

#### 3.1.2 Network hub topography

Similarly to network topology characteristic developmental differences were observed in the topography of “hubness” (BC) between newborns and infants in the two later age groups. In all three bands, AGE had a significant main effect. In the delta band, this effect was mainly due to higher BC for newborns compared to both 4 and 9 month olds. In the theta and beta bands, AGE significantly interacted with ELECTRODE. For these two bands, post-hoc tests were conducted separately for each electrode.

Within the theta network, significant AGE effect was obtained for the Fz, C3, P3, and Pz electrodes, mostly due to the significant decrease of centrality (BC) at nine months compared to birth, except for the Pz lead (Figure 3A). In the beta band, the effects of age on BC were significant at Fp1, Fp2, and P3 showing a pattern similar to the theta band (Figure 3B): BC decreases with age, except for the Fp1 lead.

**Figure 3.**
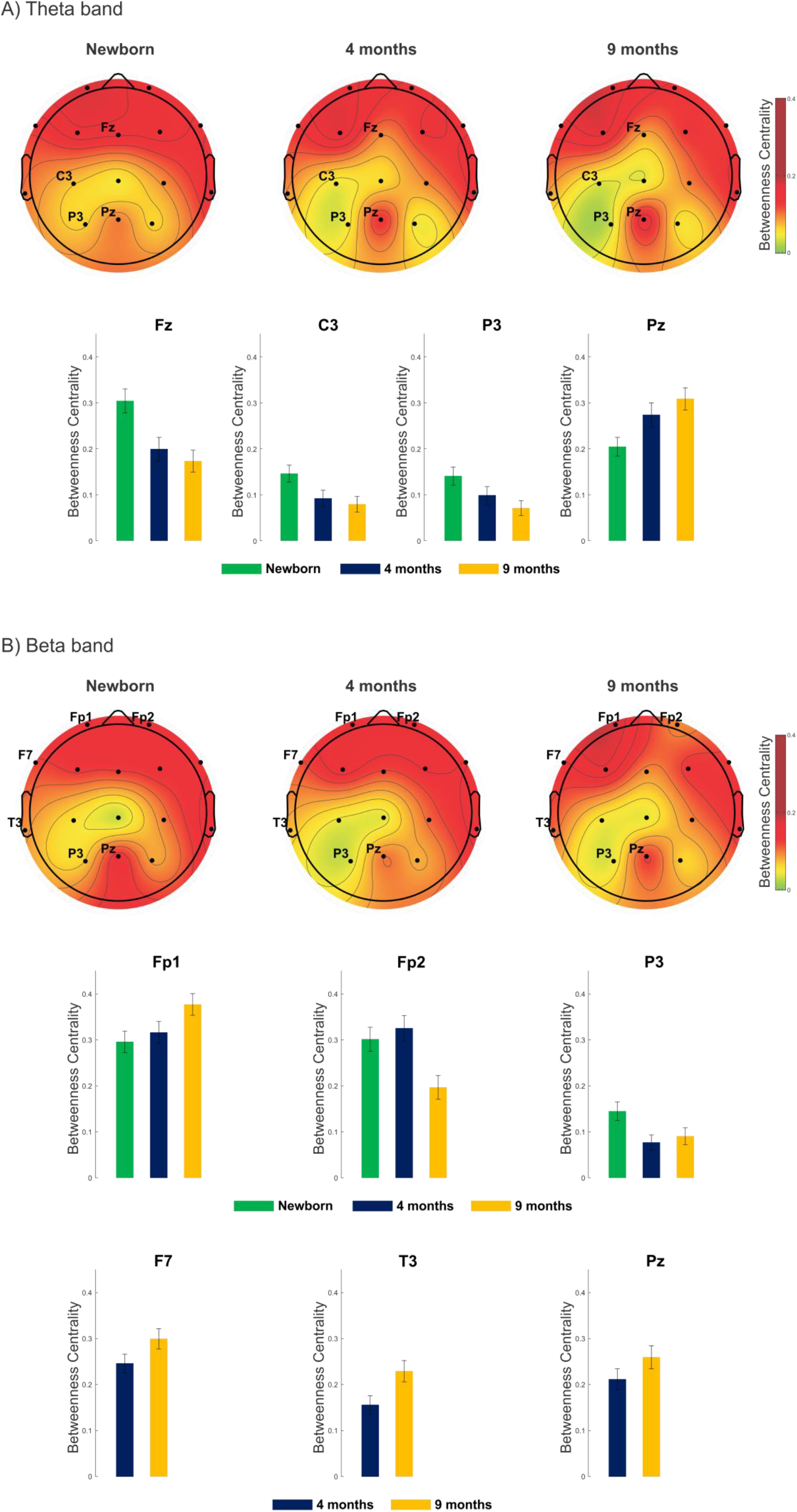
Scalp distributions for “Betweenness Centrality” (BC) in the theta (Panel A) and beta band (Panel B) at the three different ages. Significant AGE effects on BC are shown on bar charts. For the theta band (Panel A), significant effects were obtained only between newborns and the other two groups, whereas for the beta band (Panel B), significant effects were also found between four and nine month olds. The latter are shown in the bottom row of the panel. Error bars denote standard error of the mean (SEM).

Between four and nine months of age significant differences were only found in the beta band (Figure 3B). The results showed that BC at the Fp2, F7, T3, and Pz locations increased, except for the Fp2 electrode, which showed the reverse effect.

### 3.2 Development of the sensitivity for processing infant directed speech during the first year of life

Significant statistical results for the ANOVAs including all three groups are given in Table 3, while those for only the 4-month and 9-month old infants in Table 4.

**Table 3.**
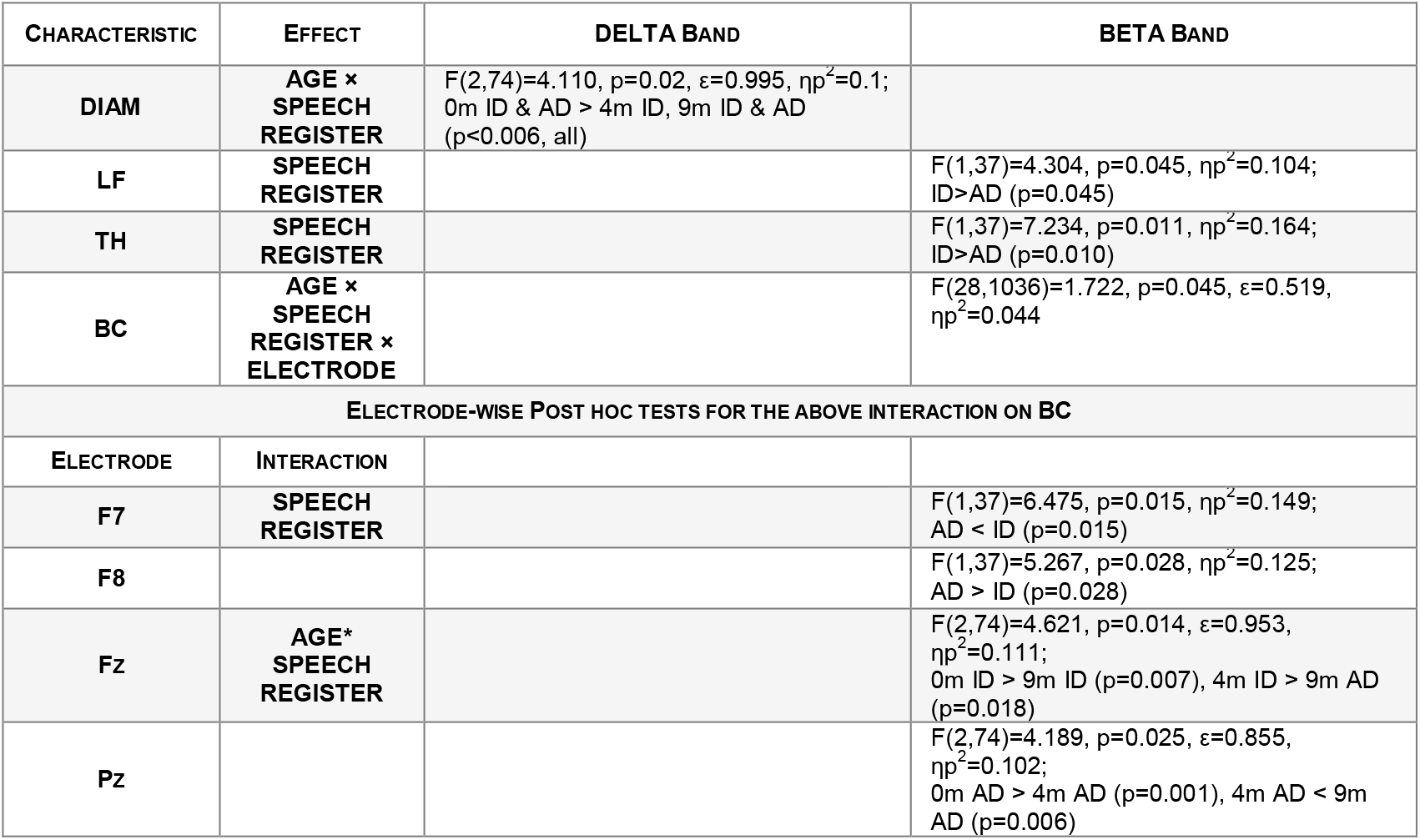
All three age groups: Summary of the significant effects involving SPEECH REGISTER on the topological (upper table half) brain network characteristics (rows), separately for each frequency band with significant effects (columns). Significant post-hoc pair-wise test results, where applicable, are shown under the ANOVA in the following format: “Age in months” [0/4/9m] “Speech Register” [ID/AD] [</>] “Age” “Speech Register” (p value). Post-hoc ANOVAs for the electrodes with significant SPEECH REGISTER effects on “Betweennes Centrality” (BC) are shown in the lower half of the table.

**Table 4.** Four and nine month olds only: Summary of the significant effects involving SPEECH REGISTER on the topological (upper table half) brain network characteristics (rows), separately for each frequency band with significant effects (columns). Significant post-hoc pair-wise test results, where applicable, are shown under the ANOVA in the following format: “Age in months” [4/9m] “Speech Register” [ID/AD] [</>] “Age” “Speech Register” (p value). Post-hoc ANOVAs for the electrodes with significant SPEECH REGISTER effects on “Betweennes Centrality” (BC) are shown in the lower half of the table.

| CHARACTERISTIC | EFFECT | THETA BAND | BETA BAND |
| --- | --- | --- | --- |
| TH | SPEECH REGISTER | | F(1,37)=4.881, p=0.033, $\eta^2$=0.117 ; ID>AD (p=0.034) |
| BC | SPEECH REGISTER × ELECTRODE | F(14,518)=2.065, p=0.04, $\epsilon$=0.567 $\eta^2$=0.053 | F(14,518)=2.823 p=0.005 $\epsilon$=0.570 $\eta^2$=0.071 |
| ELECTRODE-WISE POST HOC TESTS FOR THE ABOVE INTERACTION ON BC |  |  |  |
| ELECTRODE | INTERACTION |  |  |
| F7 | SPEECH REGISTER | | F(1,37)=6.741, p=0.013, $\eta^2$=0.154; AD < ID (p=0.014) |
| F8 | | | F(1,37)=5.628, p=0.023, $\eta^2$=0.132; AD > ID (p=0.023) |
| Fz | | F(1,37)=4.634, p=0.038, $\eta^2$=0.111; AD > ID (p=0.038) | F(1,37)=8.799, p=0.005, $\eta^2$=0.192; AD > ID (p=0.005) |
| F4 | | F(1,37)=4.478, p=0.041, $\eta^2$=0.108; AD > ID (p=0.041) | |
| C4 | | F(1,37)=6.243, p=0.017, $\eta^2$=0.144; AD > ID (p=0.017) | |
| Pz | SPEECH REGISTER × AGE | | F(1,37)=4.778, p=0.035, $\eta^2$=0.114; 4m AD < 9m AD (p=0.001) |
| C4 | | F(1,37)=4.778, p=0.035, $\eta^2$=0.114; 4m AD > 4m ID (p=0.017 ) | |

#### 3.2.1 Brain network topology

The integration capacity of the delta brain network (measured by DIAM) differentiated between the processing of speech register and that was significantly changing over the first year of infancy (AGE × SPEECH REGISTER interaction in the DIAM parameter, see Figure 4A. Specifically, DIAM within the delta band network was higher during the listening to ID relative to AD speech. The difference was separately significant at four months of age, but not at the other two measurement points, resulting in the significant interaction.

**Figure 4.**
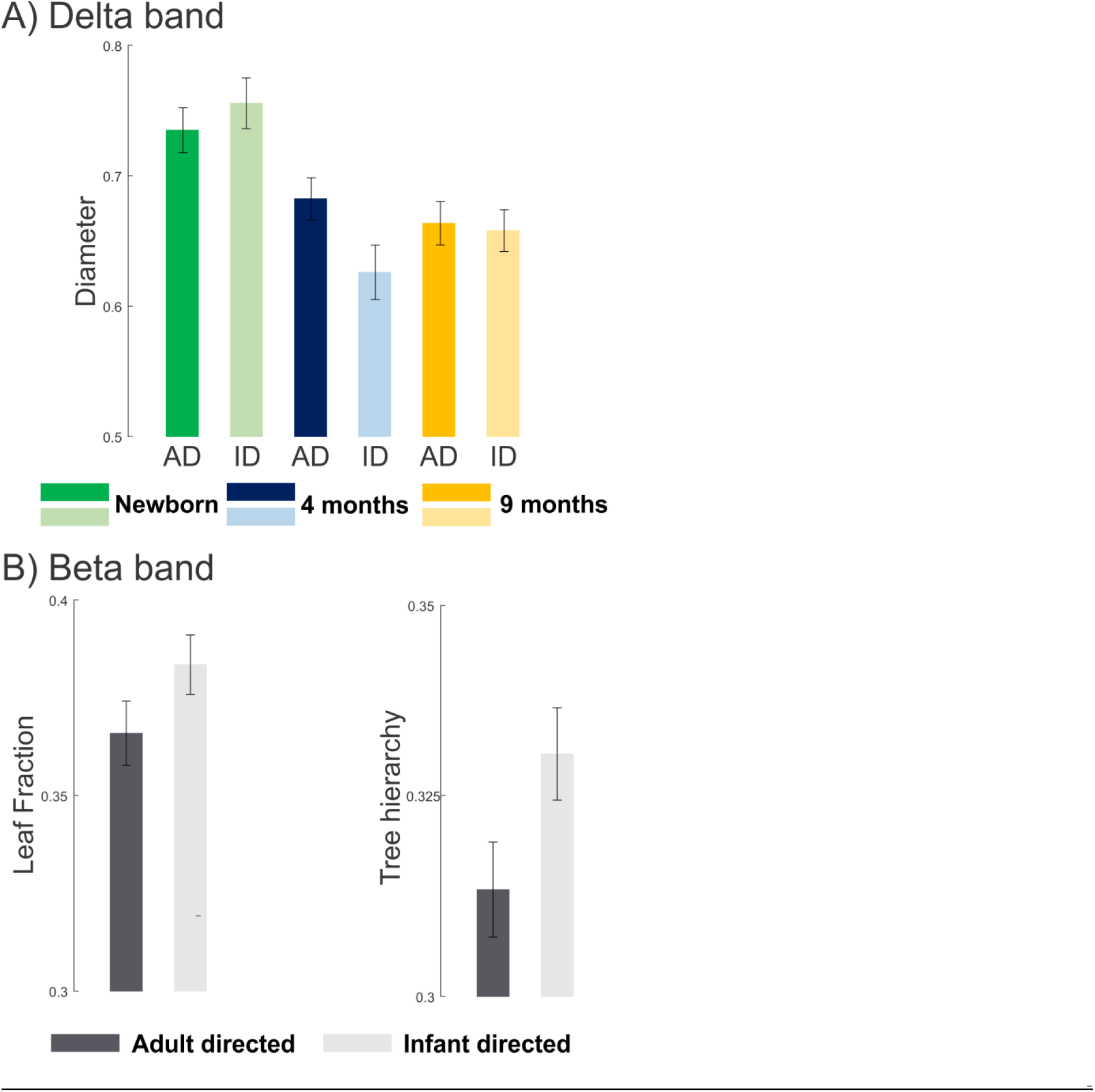
Significantly different brain network topology characteristics in processing infant vs. adult directed speech in the delta band (A) and the beta (B). A significant interaction between AGE and SPEECH REGISTER was found in the delta band, while significant SPEECH REGISTER effects were obtained in the beta band. Error bars denote standard error of the mean (SEM).

Regardless of age we observed that beta band topology differed between listening to infant-directed and adult-directed speech: communication within the beta network became from line like-like to more functionally segregated, when listening to ID relative to AD as indicated by larger Three Hierarchy (TH) and higher Leaf Fraction values for ID compared to AD (Figure 4B). When considering only 4- and 9-month-olds, this SPEECH REGISTER effect was significant only for the TH property of the beta band network (again TH showing larger value for ID compared to AD).

#### 3.2.2 Network hub topography

The scalp topography of network node “hubness” (measured by BC) differentiated the processing of speech registers and it was significantly changing over the first year of infancy, varying also across electrodes (AGE × SPEECH REGISTER × ELECTRODE interaction in BC for the beta band; see Figure 5). Post-hoc electrode-wise ANOVAs for the SPEECH REGISTER effect showed a significant difference in frontal lateralization (F7: AD<ID and F8: AD>ID) and significant AGE × SPEECH REGISTER interactions over Fz and Pz in the beta band.

**Figure 5.**
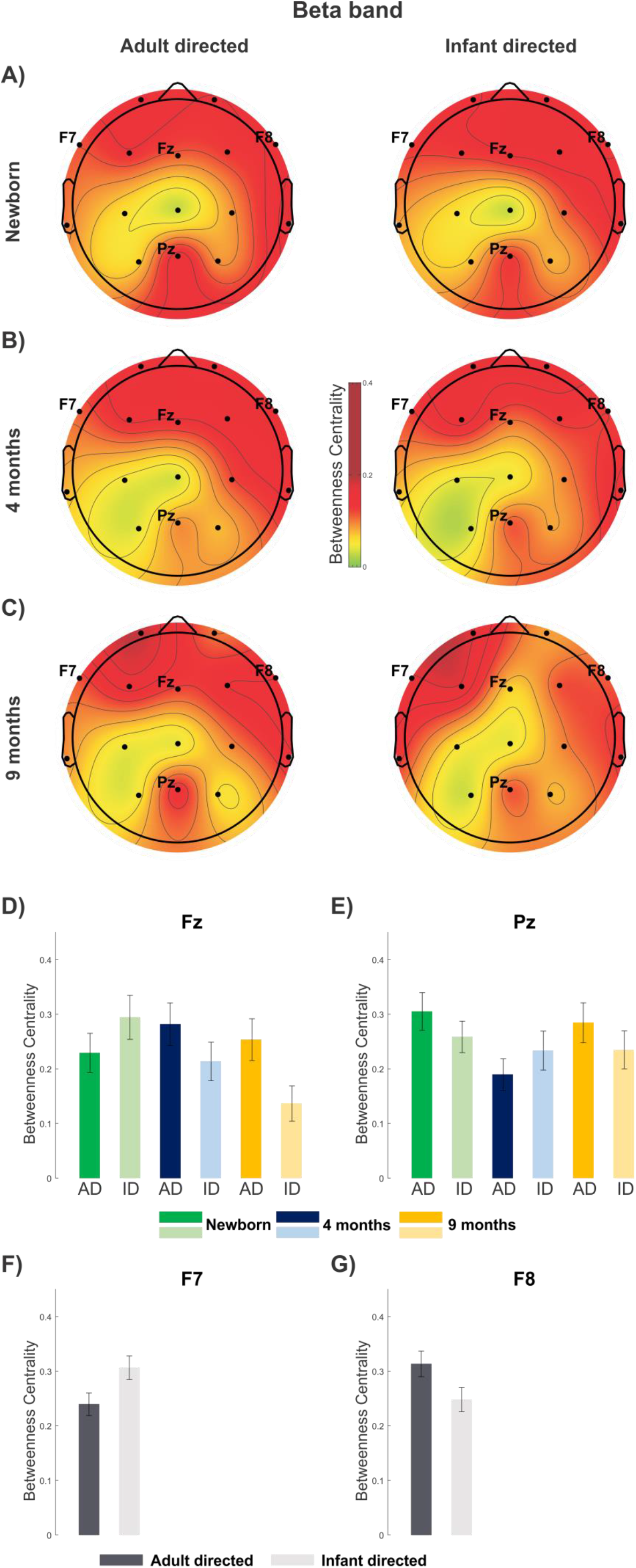
Scalp distributions for “Betweenness Centrality” (BC) in the beta band at the three different ages and speech registers (A-C) and significant AGE and SPEECH REGISTER effects on BC (D-G) with F-G only showing effects for 4 and 9 months olds. Error bars denote standard error of the mean (SEM).

When comparing BC only between 4- and 9-month old infants, the ANOVA yielded a SPEECH REGISTER main effect for electrodes F7, F8, and Fz in the beta band and for Fz, F4, and C4 in theta band (Figure 5). All of these electrodes showed higher BC for AD compared to ID speech except for F7, which showed the reversed effect. AGE × SPEECH REGISTER interactions were found in the beta band for Pz, with higher BC values for AD at 9-month compared to that at 4 months (Figure 5), and in the theta band for C4, with higher BC values for AD than ID speech at 4 months (Figure 6).

**Figure 6.**
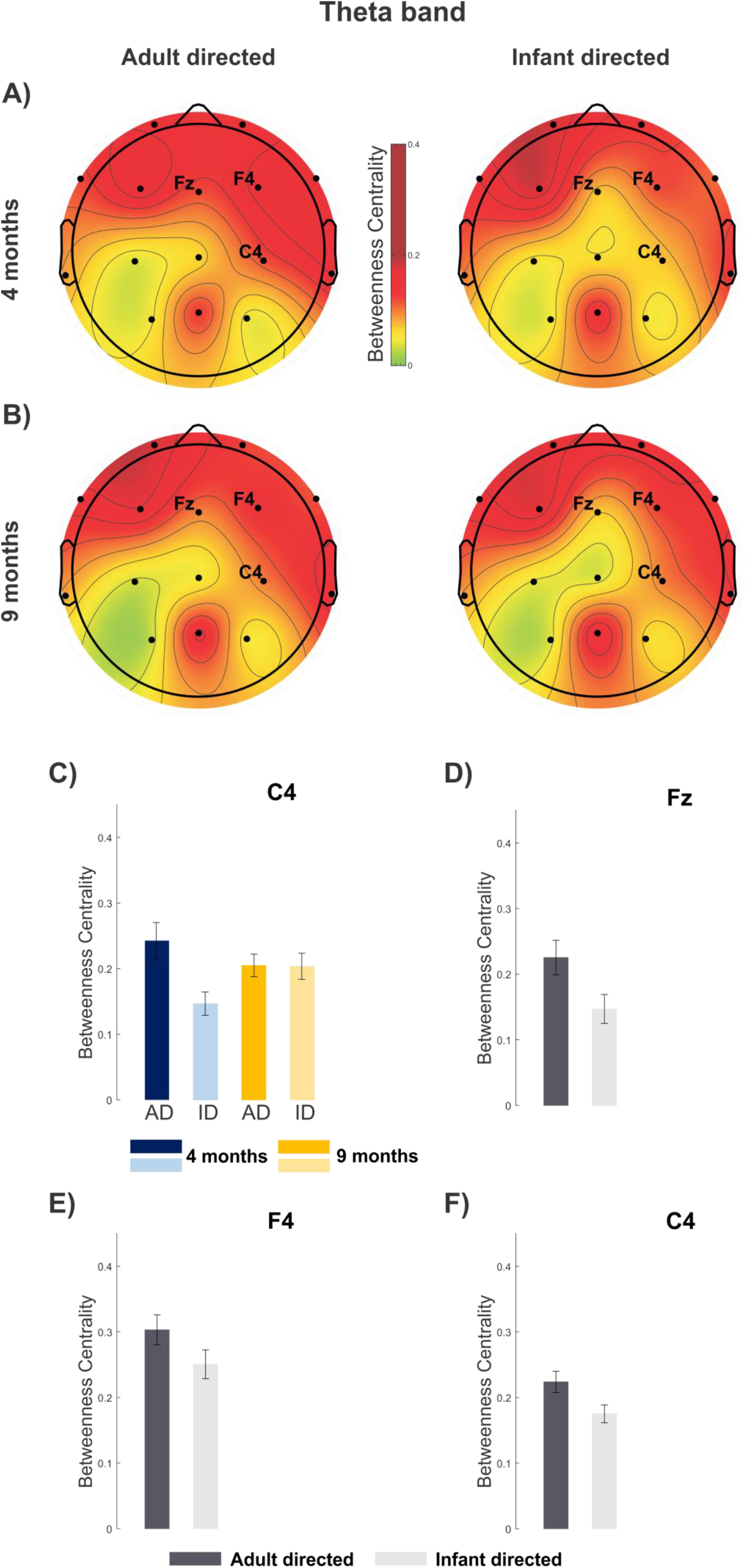
Scalp distributions for “Betweenness Centrality” (BC) in the theta band at the three different ages and speech registers (A-B) and significant AGE and SPEECH REGISTER effects on BC (D-F) Since these effects are present when only 4 and 9 months olds are considered, only those ages are represenred. Error bars denote standard error of the mean (SEM).

## 4 Discussion

Low-frequency brain oscillations in the theta and delta range synchronize to the dynamics of the speech envelope associated with syllabic (∼5 Hz) and phrasal (∼2 Hz) rates presentation respectively, while neural activity in the beta range follow the fine-grained temporal dynamics related to phonetic features (∼20 Hz; Giraud and Poeppel, 2013; Leong and Goswami, 2014; 2015). Therefore the networks operating in these frequency ranges are of general interest to speech perception and also specifically to processing IDS and ADS. In the following sections the current data will be discussed in these two contexts.

### 4.1 The development of speech processing networks during the first year of infancy

When considering the topological parameters in the delta and theta band one can discern a qualitative change in the networks between 0 months and 4 months of age. In contrast, smaller, quantitative changes can be observed between 4 and 9 months (Figure 2). This can be interpreted as a shift occurring in speech processing networks before 4 months. There are important developmental changes taking place between these ages (e.g., stronger preference for social sounds, McDonald et al., 2019; the ability to participate in peekaboo types of games, Jaffe at al., 2001; the ability to produce vowel-like sounds and babble (Kuhl, 2004)).

With regard to topographical changes in “hubness” (“betweenness centrality”, BC), in the theta band, BC decreases with age over most electrodes (except for Pz; Figure 3A). This may reflect steps towards the speech processing network observed in adults. That is, at birth there are more centralized hubs with few connections in between, whereas later the number of connections is spread across more nodes: further areas get involved in speech processing. On the other hand, the in the beta band, larger hubs over left frontal electrodes with a concurrent decrease over the right frontal ones (Figure 3B) probably reflect increasing left lateralization of speech processing, a feature corresponding to the results of several other studies (Bortfeld, Fava & Boas, 2009; Minagawa-Kawai, Cristià & Dupoux, 2011).

### 4.2 The sensitivity for processing IDS differentially changes over the first year of infancy

There was a significant effect of speech register on the topology of the brain networks active during speech in the delta and beta bands. Some difference was expected, because even newborns have been shown to process AD and ID speech differentially (Haden et al., 2020; Saito et al., 2007). The current results specified this differential sensitivity. No significant interaction was found between age and speech register on topological (network structure) measures in the beta band (Figure 4B). The beta band is associated with the processing of the “faster” aspects of speech such as phonemes (Giraud & Poeppel, 2012). The lack of age effect on the difference between the two speech registers supports the notion that infants are capable of differentiating phonemic contrasts from birth (Boesseler et al., 2016) and this function does not require large-scale structural reorganization of the underlying brain network before 9 months of age. Therefore, the large ID/AD differences are reflected by the same network structure differences from birth onwards. The higher leaf fraction and tree hierarchy network description values found for IDS across the three tested age groups point to a network that is more integrated and more hierarchical when processing infant than adult directed speech. While the leaf fraction effect is only significant when all there age groups are analyzed together, the tree hierarchy difference is present also when only 4 and 9 month olds’ data is taken into account.

The speech register effect on diameter (width) of the network in the delta band is modified by age, with only 4-month old infants showing a significant effect (Figure 4A): wider networks when listening to AD than ID speech. A possible post-hoc interpretation is that the ID speech used in the current experiments was best tuned to 4-month olds, which was the age of the actress’ own child at the time of the recording. Since characteristics of the mother’s infant-directed speech changes with the development of her child (Golnikoff et al., 2015), it is possible that 4-month old infants were better tuned to the actual ID speech presented to them than either of the other age groups. Further, the delta band is best attuned to prosodic features of speech (Leong & Goswami, 2015), and ID speech at 4 months may overemphasize prosody as at this age caretakers start to turn from the initially predominant emotional aspect of ID speech towards supporting speech segmentation (Thiessen, Hill & Saffran, 2005, which indeed develops during this period, and by 7-9 months, infants begin to extract words from speech (Kooijman et al., 2013).

The topographical properties of the network processing ID vs. AD speech are reflected in the differences in “Betweenness centrality” (BC) at different electrode locations. Higher BC corresponds to more central nodes in networks. Over frontal areas, beta-band networks show higher BC for IDS on the left and higher BC for ADS on the right, irrespective of age (Figure 5). The well-known model of brain asymmetry for language processing in adults (Friderici and Alter, 2004) posits that the left-hemispheric areas dominate the processing of syntactic and semantic information, whereas prosody receives more detailed processing in right-hemispheric areas. However an important qualifier of this account is the difference in the temporal scales over which different linguistic features are processed (Poeppel, 2003; Cogan & Poeppel, 2011). In the current data, hemispheric asymmetry arises in the beta band, which may reflect phonetic processes. Indeed, the semantic/syntactic vs. prosodic account is unlikely to explain the observed difference in lateralization. IDS vs. ADS processing shows topographical differences with regard to vowel hyper-articulation (Lovcevic et al., 2019), which is a key feature of IDS and its duration is compatible with the cycles of the oscillations producing beta-band activity. Therefore the currently observed asymmetry may reflect differences between IDS and ADS on a phonemic level. The development of hemispheric asymmetry continues well into childhood; e.g., 4-6 year old children show significant right hemispheric activation (Olulade et al., 2020). One may assume that in infants of 0-9 month age, only the initial phase of the development of the lateralization of speech processing can be observed.

Age interacted with IDS vs. ADS differences for the beta band over midline frontal and parietal electrodes. The frontal effect was mainly due to lower BC to IDS at 9 months, whereas Pz shows a dip in the BC values at 4 month olds, especially for ADS (Figure 5). Neither pattern may reflect pure speech processing. The frontal effect may fall into the general tendency of decreasing BC with age. The parietal pattern, while surprising, corresponds to a recent fMRI finding showing a “V” shaped pattern in the connectivity of language and motor areas during this period (Yin et al., 2019). The location of the apparent decrease in BC is also somewhat consistent with the involvement of motor associated areas (e.g. parts of the supramarginal gyrus, Oberhuber et al., 2016). Importantly, similarly to our study, Yin and colleagues measured the same infants at different ages. It appears that this type of interaction may remain hidden when using cross-sectional samples. Thus, again, this may, at least partly reflect a general aging effect.

Theta-band topographic effects distinguishing processing IDS and ADS are only significant for the two older age groups, and more pronounced at 4 months of age (Figure 6). Consistently with literature (Friderici and Alter, 2004; Telkemeyer et al., 2009), the right hemisphere shows increased BC values for the theta band, which has been linked with the processing of suprasegmental speech features, in general (Abrams, et al., 2008; Peelle & Davis, 2012) and in this age group (Telkemeyer et al., 2011). The midline-right localization of the effect suggests that the effect may be related to prosodic processing. However in our data, ADS produced higher BC values than IDS, while IDS typically includes more exaggerated prosodic cues. One possible explanation is that the higher BC activation for ADS stems from the larger effort necessary to process the less salient prosody of ADS.

### 4.3 Limitations of the study

An important limitation of the current study is that neonates were asleep, while 4 and 9 month olds were awake. This confound may partly explain the larger changes seen between 0 and 4 than between 4 and 9 months. (A good control would be to record 4- and 9-month old infants asleep. This, however, is not easy to do.) On the other hand, neonatal sleep is not identical to sleep later in life (Jenni, Borbély & Achermann, 2004). Some of the sleep stages stimulus processing is only altered to a smaller degree although the effect on functional network connectivity is not yet clear in infants (Taga, Watanabe & Homae, 2018).). Therefore, it is likely that at least some of the brain network differences found between newborns and the other two age groups reflect developmental changes. (Unfortunately, the recording time in newborns and the size of the group followed did not allow us to gather sufficient data for comparing between different neonatal sleep stages and the data collected at later ages.)

Another limitation is the lack of non-speech control. The validity of interpreting some of the topographical changes to the development of speech processing would have benefited from a comparison with changes occurring in general sound processing during the same period. Again, the limited recording time, especially in the two later age groups prevented the collection of control data. However, while changes towards a more distributed network may be a general tendency during early development towards the “ideal” small world networks, speech processing certainly would benefit from this tendency, and involving areas undertaking more advanced speech processing is likely to occur. On the other hand, the finding of increasing left lateralization matches well the literature. Thus it is likely specific to speech processing.

The low number of electrodes recorded (a limitation imposed by the recording equipment used for newborn data collection) precludes detailed analysis of the topography. Given the low temporal resolution and the ambiguity of the sources our analysis was limited to lateralization and inference to gross brain areas along the frontal-central-parietal direction, which are sufficiently reliable in the data. Further, the topological measures, while also less than ideally detailed, reflect important large-scale structural properties of the networks (Tóth et al., 2017).

The caretakers’ infant-directed speech changes together with the development of their child (Englund & Behne, 2006 however see also, Kalashnikova & Burnham, 2018). Thus it is a difficult choice, whether to present the same ID stimulus to different age groups (which allows comparability in the acoustic/phonetic sense) or different, age-appropriate ID speech segments (which may better match the infants’ current experience with ID speech). We chose the first alternative, but acknowledge that some effects could have been stronger by the second alternative. A future study could contrast these two strategies within the same infants.

## 5 Conclusions

The general picture emerging from the results is that with development the speech processing network becomes more integrated and its focus is shifting towards the left hemisphere. However, development is far from a linear increase in some properties. The network is reorganized in multiple stages of maturation with different aspects of speech processing, as reflected by different EEG bands, occurring in parallel, but with different timing. The role of IDS changes during the first year of life, probably accompanied by changing properties of IDS itself. As a result, the various features of IDS may become more or less important at a given time, which affects sensitivity to IDS rise and fall in different networks sub-serving speech processing. The pattern of results obtained in the current study describes various details of these developmental changes in the first nine months of life.

## Acknowledgement

The authors are grateful to Eszter Lányi for help with figures and Kinga Kerner for an earlier version of the data analysis. The authors are also grateful to the staff of RCNS (Judit Roschéné Farkas, Lívia Özéné Kende, Zsófia Haraszti) who recorded the data and finally to all participants and their families.

## Notes

**Declarations of interest** The authors declare no conflicts of interest.

**Funding** This work was funded by the Hungarian National Research Development and Innovation Office (ANN131305 and PD123790 to BT, and K115385 to IW); the János Bolyai Research Grant awarded to BT (BO/00237/19/2) and separately to GPH (BO/00523/21/2) and the New National Excellence Program of the Ministry for Innovation and Technology from the source of the National Research, Development and Innovation (ÚNKP-21-5-BME-364) for GPH. This work was supported by project no. K115385 (all authors) and ANN131305 (BT) that has been implemented with the support provided by the Ministry of Innovation and Technology of Hungary from the National Research, Development and Innovation Fund, financed under the NKFI K and NKFI ANN funding scheme (respectively).

https://osf.io/kd2tp/?view_only=5988023d9377426382210675b7170ac9

### Competing Interest Statement

The authors have declared no competing interest.

